# Unravelling processes between phenotypic plasticity and population dynamics in migratory birds

**DOI:** 10.1101/2021.02.15.429667

**Authors:** Jin Liu, Weipan Lei, Xunqiang Mo, Chris J. Hassell, Zhengwang Zhang, Tim Coulson

## Abstract

1. Populations can rapidly respond to environmental change via adaptive phenotypic plasticity, which can also modify interactions between individuals and their environment, affecting population dynamics. Bird migration is a highly plastic resource-tracking strategy in seasonal environments. However, the link between the population dynamics of migratory birds and migration strategy plasticity is not well understood.
2. The quality of staging habitats affects individuals’ migration timing and energy budgets in the course of migration, and can consequently affect individuals’ breeding and overwintering performance, and impact population dynamics. Given staging habitats being lost in many parts of the world, our goal is to investigate responses of individual migration strategies and population dynamics in the face of loss of staging habitat, and to identify the key processes connecting them.
3. We started by constructed and analysed a general full-annual-cycle individual-based model with a stylized migratory population to generate hypotheses on how changes in the size of staging habitat might drive changes in individual stopover duration and population dynamics, and to identify the key processes connecting them. Next, through the interrogation of census data, we tested these hypotheses by analysing population trends and stopover duration of migratory waterbirds experiencing loss of staging habitat.
4. We found empirical support for our modelling-identified hypotheses: the loss of staging habitat generates plasticity in migration strategies, with individuals remaining on the staging habitat for longer to obtain food due to a reduction in per capita food availability. The subsequent increasing population density on the staging habitat has knock on effects on population dynamics in the breeding and overwintering stage.
5. Our results demonstrate how environmental change that impacts one energetically costly life history stage in migratory birds can have population dynamics impacts across the entire annual cycle via phenotypic plasticity.

## Introduction

Populations can rapidly respond to environmental change via adaptive phenotypic plasticity, and this allows them to cope with profound environmental impacts (Pigliucci 2001, Piersma and Drent 2003, Coulson et al. 2017). Plasticity modifies interactions between individuals and their environment, ultimately affecting population dynamics (Miner et al. 2005). Migration can be an adaptive plastic strategy in seasonal environments (Lack 1968, Newton 2010), that allows individuals to increase reproductive output by avoiding unsuitable ecological conditions (Hedenström 2008, Winkler et al. 2014). Plasticity of migration strategies enables migratory species to respond to environmental changes in multiple ways, such as by altering migratory routes (Sutherland and Crockford 1993, Dolman and Sutherland 1995), timing of migration (Gienapp et al. 2007, Balbontín et al. 2009), and through diet (Parrish 2000). However, the link between the population dynamics of migratory species and migration strategy plasticity is not well understood.

Bird migration is a resource-tracking strategy that aims to optimize a bird’s energy budget in the face of fluctuating resources in seasonal environments, and in the face of strong competition (Cox 1968, Alerstam et al. 2003, Somveille et al. 2018, Winger et al. 2019). Migration is energetically costly, so birds build up fat reserves. However, carrying a large energy reserve increases flight costs and can also attract predators (Alerstam and Lindström 1990). One strategy to minimise such costs is to stop over several times during the journeys between breeding and wintering sites to refuel (Piersma 1987). For individuals to remain in favourable environments across their migration route, they must carefully manage the timing of departure and arrival (Alerstam and Lindström 1990, Alerstam et al. 2003, Winkler et al. 2014). In general, individuals that arrive at breeding grounds earlier have higher reproductive success than those that arrive later (Marra et al. 1998, Norris et al. 2004), and selection favours individuals that minimize the time spent travelling during the northward migration (Lindstrom and Alerstam 1992). Migratory birds usually spend much longer accumulating energy reserves in staging areas than in flying (Hedenström and Alerstam 1997). Therefore, the total time spent on migration is consequently strongly influenced by the quality of, and an individual’s behaviour at, staging areas (Hedenström and Alerstam 1997, Erni et al. 2002).

For migratory species, all stages of the annual cycle are closely linked at both the individual- and population-level, through carry-over and density-dependent effects (Newton 2010, Harrison et al. 2011). The individual state in one stage can influence individual performance in subsequent stages, and the change in population size in one stage can influence per capita rates and consequently regulate population size in later stages (Marra et al. 1998, Studds and Marra 2005, Ryan Norris and Marra 2007, Ratikainen et al. 2008). The strategy an individual follows while at the staging area can consequently affect breeding and overwintering performance, and impact population dynamics.

Staging habitat for migratory waterbirds in the Yellow Sea is being lost in significant quantities, primarily due to land reclamation for infrastructure development and aquaculture (Yang et al. 2011, Bi et al. 2012, Murray et al. 2014, Wang et al. 2014). It has been suggested as the main contribution to population declines of migratory birds along the East Asian Australasian Flyway (EAAF) (Amano et al. 2012, Ma et al. 2014, Piersma et al. 2016, Studds et al. 2017), presumably because staging habitat in this system is the stage of the annual cycle where density-dependence is strongest (Sutherland 1996b). Habitat loss in staging habitats can reduce food resources, decrease foraging and fat-accumulation rates of migrants (Baker et al. 2004, Morrision 2006, Verkuil et al. 2012), increase competition and interference in the population, and can have significant consequences for population regulation (Sutherland 1996b, Newton 2010). However, the way in which individuals respond to such changes, as well as the processes and mechanisms that cause population declines, are yet to be generally established. Migratory waterbirds along the EAAF provides a unique system to explore how individuals respond to changes in one life history stage, and how this response would influence their populations.

In this study, we use theoretical modelling to generate hypotheses that we next test with empirical data to examine the effects of habitat loss within staging areas on individual migration strategies and population dynamics, and the process that connect them. First, we conducted an individual-based modelling (IBM) exercise, building a stylized full-annual-cycle model in which individuals follow the same migrating rules, to investigate how variation in the size of staging habitat drives changes on individual stopover duration and energy reserves, population density, and affects life-history process including reproduction and survival at different life cycle stages. We used this model to make predictions about changes in strategy might influence population dynamics in the staging, breeding, and overwintering grounds. We did this by examined where in the life cycle density-dependence operated most strongly. Next, using 13 years of census data, we examined whether observed empirical trends were consistent with hypotheses we generated from our individual-based model. Our empirical analyses were consistent with the theoretical hypotheses our model suggested: we found that the loss of staging habitat generates plasticity in migration strategy, with individuals remaining in the staging habitat for longer to obtain food due to a reduction in per capita food availability. The subsequent increasing population density in the staging habitat has knock on effects on breeding and overwintering stage that impact the population dynamics, via modified survival and reproduction rates. We conclude that environmental change effects on one life history stage in migratory birds can consequently have population dynamics impacts across the entire annual cycle via phenotypic plasticity.

## Materials and Methods

### The individual-based model (IBM)

The IBM we constructed is a stylized model. The basic assumptions of our model are:

1. the stages of the annual cycle in our model includes northward migration, staging in the course of northward migration, breeding, southward migration, and overwintering;
2. all individuals followed the same set of rules in our models; 3) individuals who meet the condition for reproduction produce once each year; 4) males are not limiting and can be ignored such that we construct a female-only model.

#### Model description

Our IBMs includes three types of habitats in the model landscape, which are wintering habitat, breeding habitat and staging habitat. The staging habitat split into two habitat types – S1 and S2 area. In order to simulate the process of habitat loss in the staging habitat, the size of S1 area remained constant across all simulations, while the habitat size of S2 area was adjusted (Fig.S1a-c). The rest of the grid cells in the landscape are non-habitat, where individuals pass by during migration but do not stop. Each grid cell within those three types of habitats contained renewing food resources, food resources renewed each time step at the habitat-specified food recovery rate after consumption. Each time step in this model represented one day such that one year was comprised of 365 steps.

We only considered the female component of the population and characterised each individual by identity, age, reproduction status, and energy reserves. The behaviours of each individual in the model included: fly, move, search for food, eat, orient, mature, reproduce and die. Fly was the movement across habitats during the course of migration, the speed of “fly” was three grid cells per time step. Move was the movement within each habitats during the course of overwintering, staging and breeding, the speed of “move” was one grid cell per time step. Birds lost energy reserves through either type of movement in our model. Birds searched for food by following a random movement rule, towards the grid cell with the highest food value, ate and increased their energy reserves when food was available, but there was no randomness of food acquisition, as long as the food resources was available, the behaviour of “eat” happened and energy was stored. When the energy reserve of an individual reached the energy threshold for departure, or the time reached for the latest possible departure arrived, the individual first oriented, adjusting its facing to the central grid cell of the next destination habitat in the next time step, then flied towards it. Individuals whose energy reserves reached the threshold for reproduction got matured and reproduced once each year. The order of “mature” and “fly to the breeding habitat” is different in the models with different breeding strategies (see below). Reproduction also consumed individuals’ energy reserves. Newborns were set as an initial value of age, reproduction status, and energy reserves. Individuals aged 15, or with zero energy reserves, died and were removed from the population. Details of events and decisions of the models are provided in Appendix S2 (supporting information).

The energy reserves of each individual was assumed to be dependent on their initial energy, energy gained, and energy expended. The expected energy gained from food relied on both population density and food density (Goss-Custard et al. 2002),. The stopover duration of a bird in the staging habitat was related to the energy requirement for migration, and the rate of energy acquisition (Hedenström and Alerstam 1997). In our model, the conditions for leaving the staging habitat were either the energy reserves exceeded the energy threshold or time passed the time threshold. Parameter values are provided in Table S1 and Table S2 in Appendix1, details of individual energy reserves and stopover duration are provided in Appendix S2 (supporting information).

#### Model implementation

Our main goal is to examine the effects of habitat change in the staging habitat on individual stopover duration in the course of northward migration and population dynamics across the entire life cycle, and to identify the processes that connect them. To examine the role of the staging in the course of northward migration across the whole life cycle to the hypothesized processes, we test impacts of reducing carrying capacity at other life cycle stages (breeding stage and wintering stage). In addition, since individuals with different breeding strategies (capital breeding and income breeding) has different energy budgets along the life cycle, we also tested effects of breeding strategies on model outputs.

Five types of models were constructed in our study, with models differing in the size of the S2 area, the size of the breeding habitat and the size of the wintering habitat, the food recovery rate in each of these three habitats, and the breeding strategies individuals followed (Fig S1, Table S1 in Appendix S1). More specifically:

- Model 1 is the null model, with the same size and the food recovery rate at all three habitats, and a “capital breeding” strategy for reproduction.
- Model 2 examined the effects of the size of the staging habitat on individual strategies and population dynamics, with eight scenarios where the size of S2 area was varied were run, with the lowest food recovery rate at the staging habitat and a “capital breeding” strategy for reproduction.
- Model 3 examined effects of the lowest carrying capacity at the breeding habitat to individual strategies and population dynamics, with the lowest size and food recovery rate at the breeding habitat and a “capital breeding” strategy for reproduction.
- Model 4 examined effects of the lowest carrying capacity at the wintering habitat to individual strategies and population dynamics, with the lowest size and food recovery rate at the wintering habitat and a “capital breeding” strategy for reproduction.
- Model 5 examine the effects of breeding strategies to individual stopover strategies and population dynamics, with the same settings of habitat size and food recovery rate with model 2, but with an “income breeding” strategy for reproduction.

The average daily population density and the total number of individuals, individual stopover duration, individual energy reserves and energy accumulation rate during stopover, were recorded at equilibrium, and were examined by comparing the mean value of simulation results. The model was run 10 times for each scenario, and for 30 years in each simulation, which was sufficient to converge to stationary dynamics. Results were obtained from year five to year 30 of the simulation (Fig S3, Appendix S1).

The average daily population density was the average of daily bird number in the habitat during the period of the current life cycle stage each year, it was recorded at the S1 area. The total number of individuals was recorded for the population and three types of habitats each year respectively. Individuals were recorded in age classes and reproduction status (juveniles, breeding adults, nonbreeding adult).

The per capita reproduction rate for adults, the reproduction rate among breeding adults and the survival rate were all calculated. The per capita reproduction rate was the ratio of the number of juveniles and the number of all adults in the breeding stage; the reproduction rate among breeding adults was the ratio of the number of juveniles and the number of breeding adults; survival rate was split into two periods, one is the survival rate of the northward migration (the ratio between the number of adults at the breeding habitat and the number of individuals on the first day of each year), one is the survival rate of the southward migration (the ratio between the number of individuals at the wintering habitat and the number of individuals at the breeding habitat).

Mean individual stopover duration in the S1 area was recorded each year. Individual departure energy reserves was recorded for three types of habitats respectively, as the energy reserves at the last day before the individual left the habitat range. Individual energy reserves during the stopover period was recorded at each time step when the individual stayed in the staging habitat. The energy accumulation rate during stopover was calculated by dividing energy gained from the staging habitat by stopover duration.

#### Sensitivity analysis

To examine the impacts of parameter values on the outputs of model 2 and to test the robustness of the model results were to variation in parameter values, we conducted a local sensitivity analysis (methods and results see Appendix S3 in the supporting information).

### Empirical data

Our empirical study was conducted in the wetlands in the north of Bohai Bay, between 38°36’-39°13’N and 117°11’-118.22’E, located in the north-west of the Yellow Sea. Census data of migratory waterbirds were collected at boreal spring between 2004 and 2018 (details of study area and data collection are provided in Appendix S5, supporting information).

#### Statistical analyses for bird number trends

The abundance of all waterbirds, the abundance of the most common species, and waterbird species richness were analyzed in this study. Survey sites that were surveyed on less than 30% of survey dates were excluded from the analysis.

To estimate total waterbird abundance, we summed the counts of all species *i* (including unidentified species) observed from all of the survey sites *k* in each day *j*, denoting it as *N*_*a*_. To identify the most common species, we used two methods: the “Frequency Based Method” and the “Distribution Based Method” to select 25 common species (Appendix S5). The number of these 25 species observed on each day *j* and survey site *k* was denoted as *N*_*b*_. To estimate waterbird richness, we calculated the number of species observed each day *j* each survey site *k*, excluding unidentified species, denoting it as *M*.

We calculated survey effort as the number of observers each day at each site as *E*_*j,k*_. We transformed date to Julian date *t*, and calculated *t*^*2*^ as visual examination of the data revealed a quadratic relationship. We treated ‘year’ (*y*) as a continuous variable in our models, and we treated ‘survey site’ (*k*) as a categorical variable.

Waterbird abundance and richness (including *N*_*a*_, *N*_*b*_ *and M*) were used as response variables, *E*_*j,k*_, *k, t, t*^*2*^, *i* and *y* were explanatory variables. The distribution of *N*_*a*_, *N*_*b*_ and *M* was well described as an over-dispersed Poisson (variance greater than mean), so we fitted Generalized Linear model (GLM) with a “quasi-Poisson” error structure in program R. The regression equations were of the form:

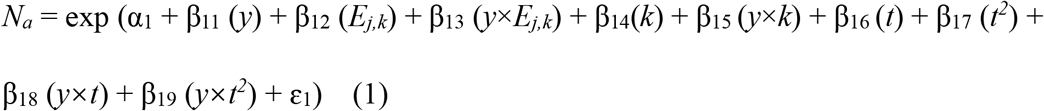

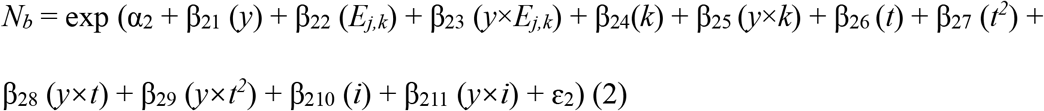

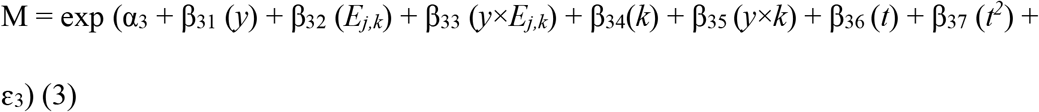

#### Statistical analyses for stopover duration trends

We estimated the stopover duration for each common species by using our census data. Since the quadratic relationship between Julian date and bird abundance revealed from our census data (Fig. S4 a), we first estimated normal distributions of bird abundance within the period of stopover for each species each year (Fig. S4b). To do this, we extracted the mean date and variance in date for each species each year from census data, and scaled the curves by the bird number from the census data. Then we estimated the date by which each quantile of the distribution of bird abundance is reached, date at 2.5% quantile and 97.5% quantile was the date of arrival at staging habitat and the date of departure from staging habitat for each species each year within 95% confidential interval respectively (Fig. S4c). We calculated the stopover duration based on the arrival date and departure date for each species each year, denoting it as *T*_*i,y*_ (details are provided in S5.4 Appendix S5).

The distribution of *T*_*i,y*_ was normal distribution, so we fitted Linear model in program R to test the stopover duration trends across years for each common species, the regression equation was of the form:

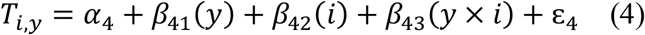

#### The correlation between stopover duration and bird abundance

We fitted a regression between bird abundance and stopover duration to test the correlation between them. We calculated the mean abundance during the stopover period for each species *i* each year *y*, denoting it as *N*_*c*._ *N*_*c*_ was used as the response variable, *y, i, T*_*i,y*_ were explanatory variables. The distribution of *N*_*c*_ was an over-dispersed Poisson, so we fitted GLM with a “quasi-Poisson” error structure in program R. The regression equations were of the form:

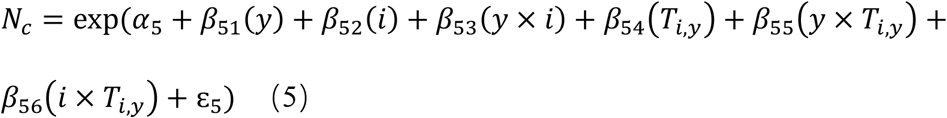

We used the ANOVA command in R to assess the significance of each variable, used adjusted-R^2^ to assess the goodness of fit of the linear model, and used 1- (Residual Deviance/Null Deviance) to assess the goodness of fit of the GLMs.

## Results

### IBM: population dynamics and individual stopover duration at the staging habitat

As the size of S2 area decreased through eight scenarios in the model 2 simulations, there was a decrease in the total number of individuals, and an increase in average daily population density at the S1 area (Fig. 1a). The curve of daily population density at the S1 area became steeper in the early and late phases of the stopover period, and higher and wider when the population reached peak numbers (Fig 1b), showing that the population took less time to reach the peak number and remained at the peak number for a longer period of time. Individual’s annual stopover duration at the S1 area increased (Fig 1c), suggesting individuals stayed longer in the S1 area as the size of S2 decreased.

**Figure 1.**
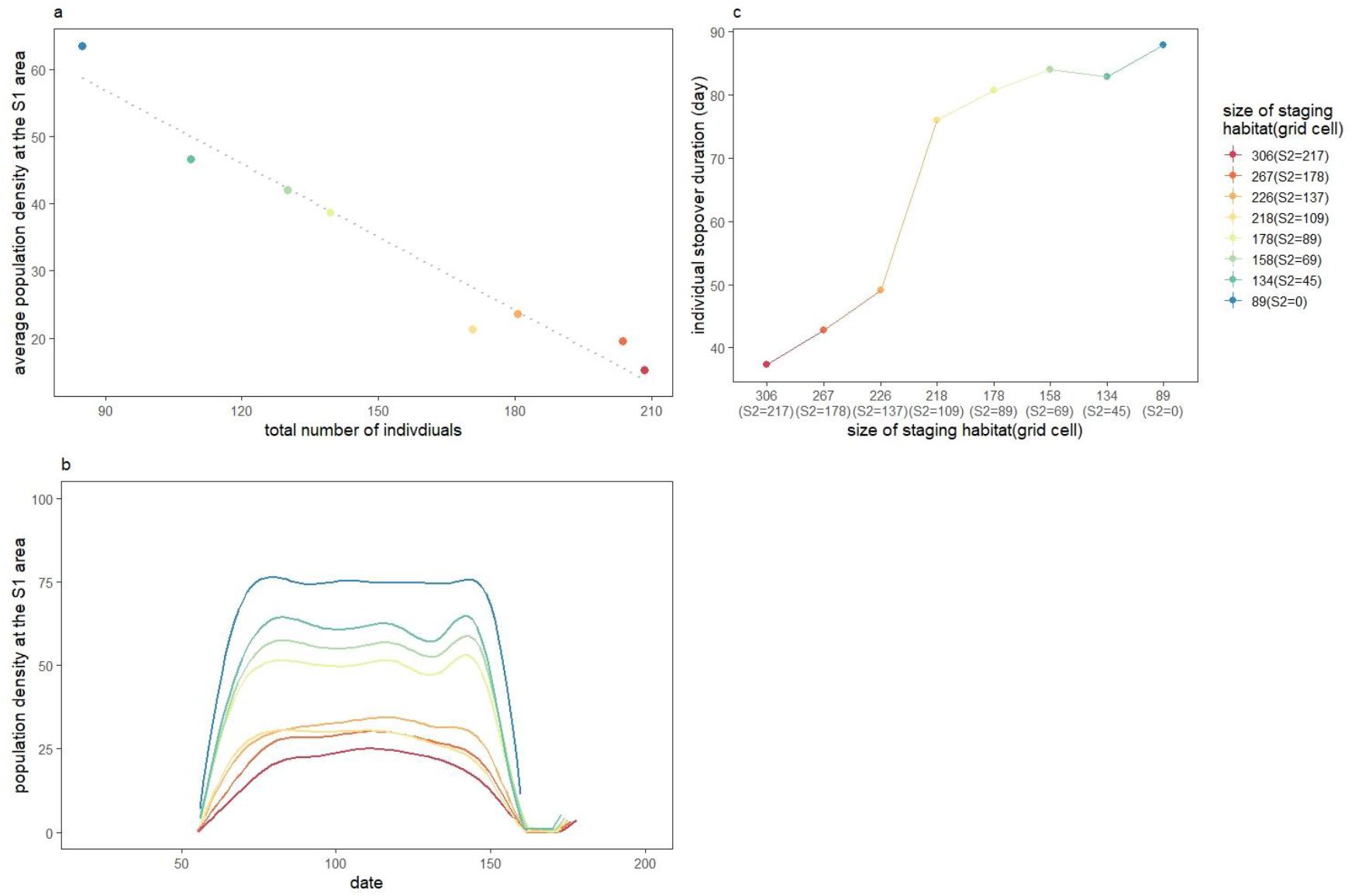
Population dynamics and stopover duration as the size of the staging habitat decreased across the eight scenarios in model 2 simulations. a) The relationship between average daily population density at the S1 area and the total number of individuals; b) The curves of population density at the S1 area during the period of stopover; c) Individual annual stopover duration in relation to the size of staging habitat.

### IBM: Individual energy reserves

As the size of S2 area decreased in model 2 simulations, the individual energy accumulation rate when birds remained in the staging habitat decreased (Fig 2a), the distribution shifted towards lower energy reserves with fewer individuals having reached the energy threshold for departure from the staging habitat (Fig 2b). The energy reserves when individuals left the staging habitat was decreased, however, the departure energy reserves from the breeding habitat and wintering habitat was increased, both in adults and juveniles (Fig 2c).

**Figure 2.**
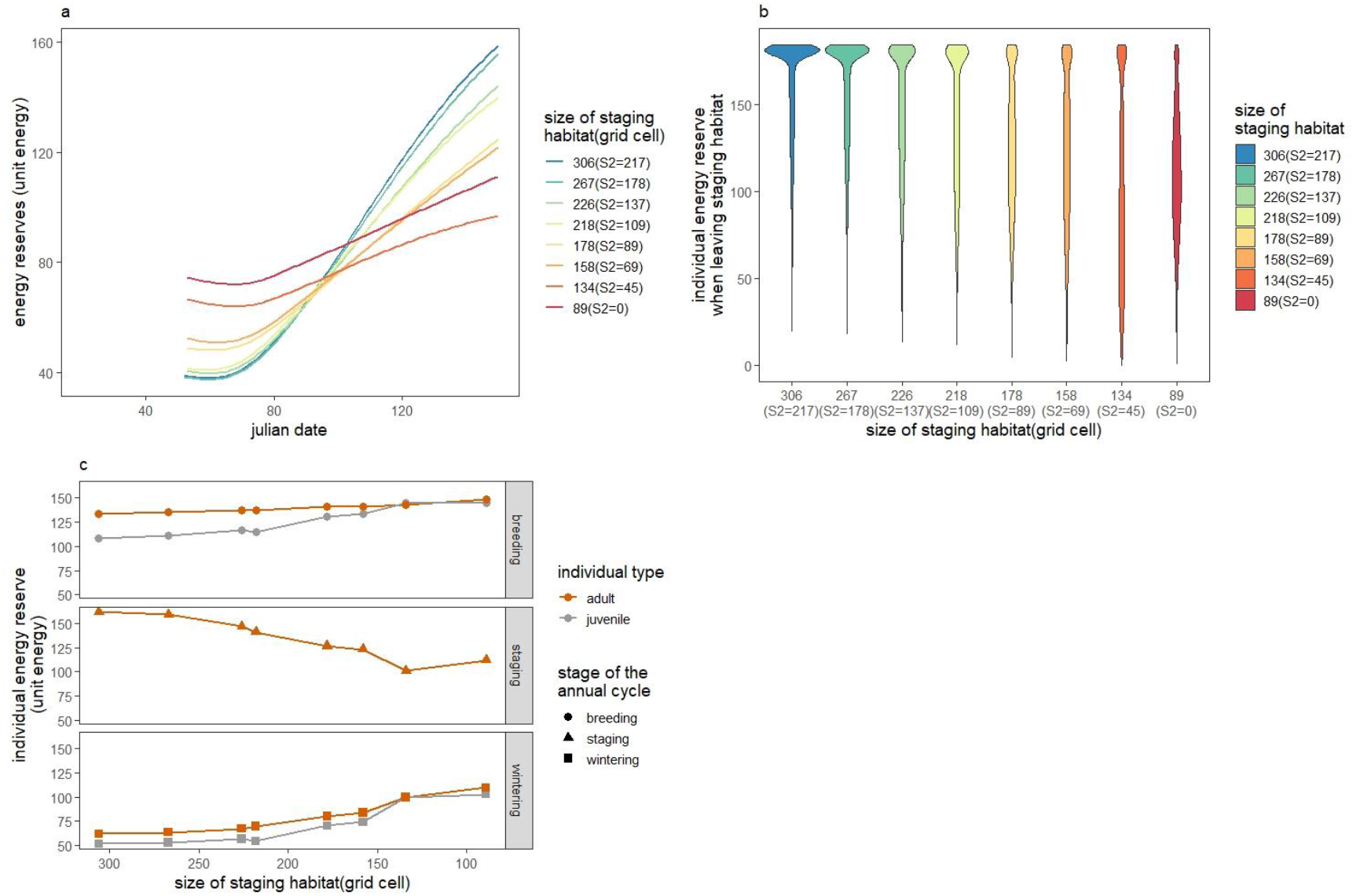
Individual energy reserves across the annual cycle in model 2 simulations. a) The energy accumulating rate during the period of stopover when individuals at the staging habitat. b) The distribution of individual departure energy reserves from the staging habitat. c) Individual departure energy reserves at each annual cycle stage.

### IBM: Population structure and demography

As the size of S2 area decreased in model 2 simulations, the proportion of breeding adults and juveniles in the population decreased significantly, while the proportion of nonbreeding adults increased in each life cycle stage (Fig 3a). The per capita reproduction rate decreased because of an increased proportion of nonbreeding adults, but the reproduction rate among breeding adults increased. The survival rate in both the northward migration and southward migration increased (Fig 3b).

**Figure 3.**
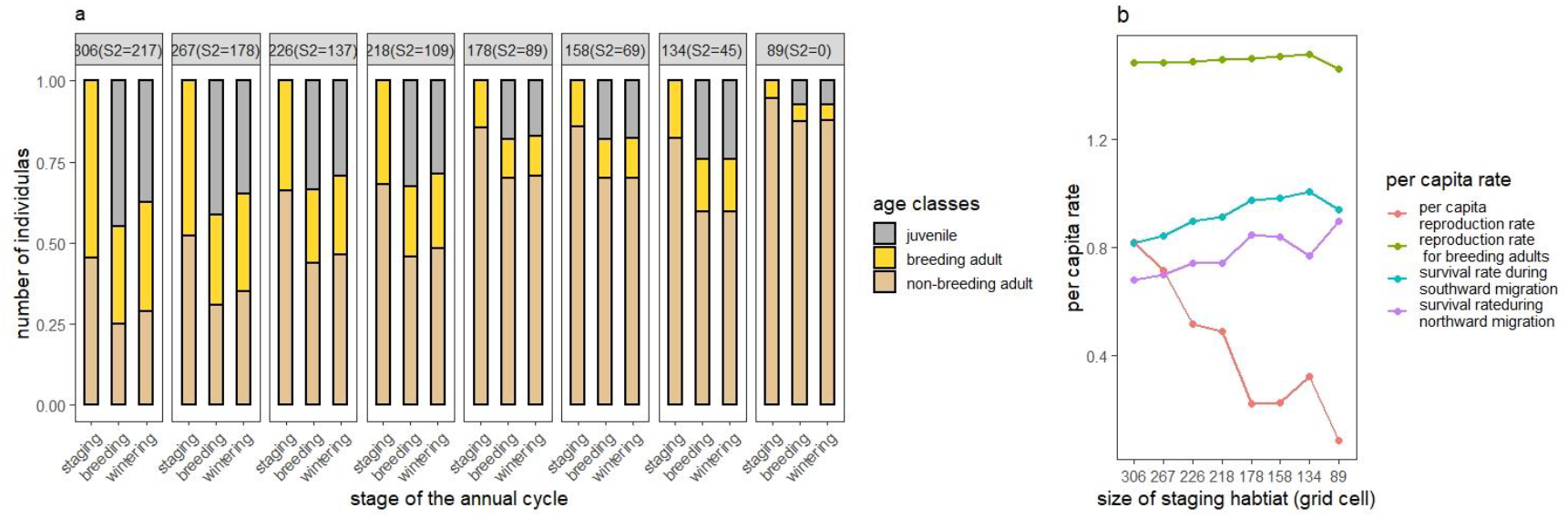
Population structure and demography as the size of the staging habitat decreased across the eight scenarios in model 2 simulations. a) The proportion of individuals in different age groups and maturity status in three stages of the annual cycle. b) per capita rates of reproduction for all adults, reproduction for breeding adults, survival during southward migration and survival during northward migration.

### IBM: the comparison between different model settings

By comparing the results from models (model 2, 3, 4) with the lowest carrying capacity at different life cycle stages and the results from the null model (model 1), we found only when the lowest carrying capacity was at the staging habitat (model 2), population density at the S1 area increased, individuals remained longer at the staging habitat, with lower energy reserves when leaving the staging habitat, leading to fewer breeding adults and juveniles in the population and lower per capita reproduction rate. Individuals do breed had better performance comparing to other models, with higher energy reserves when leaving the breeding and wintering habitat, and higher survival rate during migration. However, when the lowest carrying capacity was at other life cycle stages (model 3 - breeding stage or model 4 - wintering stage), the processes was completely different. By comparing the results from models with different breeding strategy settings (model 2 – capital breeding, model 5 – income breeding), we found different breeding strategies did not influence the processes linking migration strategy and population dynamics across the life cycle (Fig 4, details are provided in Appendix S4).

**Figure 4.**
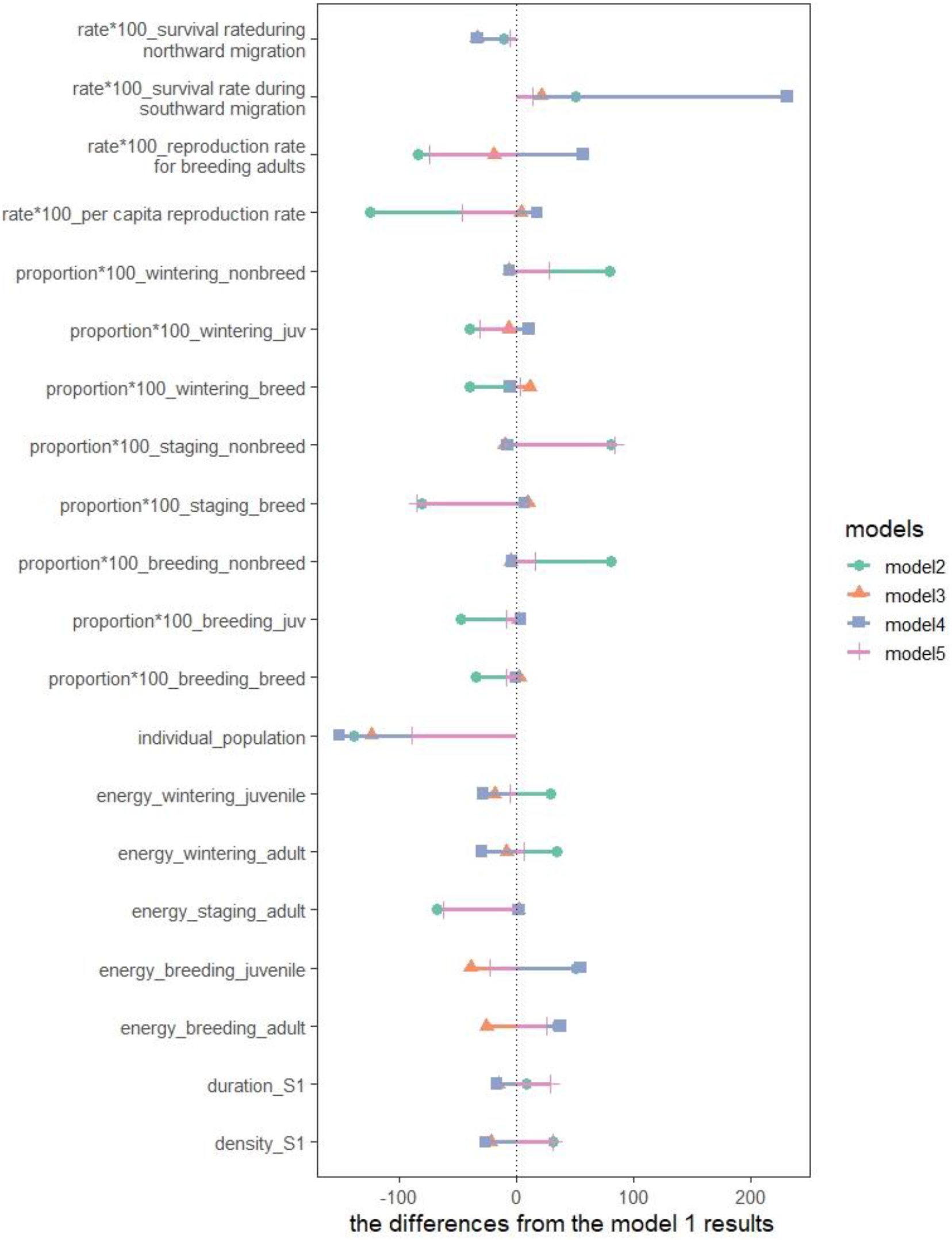
Comparison of models outputs. The name of results are in the form of “the type of results_habitat type (or the whole population)_age classes”. Six types of results are listed, “density” is the average daily population density, “duration” is the individual stopover duration, “energy” is the energy reserves when leaving the habitat, “individual” is the number of individuals, “proportion^*^100” is the one hundred fold proportion of individuals of each age class in the population, “rat*100” is the one hundred fold vital rates. Four types of habitat are listed, including the S1 area, the staging habitat, the breeding habitat, and the wintering habitat. Age classes includes juveniles and adults, adults were split into two types: breeding adults and non-breeding adults. The vertical black dotted line represents the position of the zero value. The distance from the point to the zero value line represents the difference between the results of the model (model 2-5) and the results of the null model (model 1).

### Empirical data: overall abundance

Analysis of the abundance of all recorded species revealed a significant increase (Fig.5a) in numbers of waterbirds at a rate of 61.85% per year (F_1, 809_ = 13.60, p < 0.001). Even though effort significantly affected abundance estimates (F_1, 808_ = 11.04, p < 0.001) with more birds counted as effort increased; there was no interaction between year and effort (F_1, 787_ = 0.094, p = 0.76). There was a positive relationship between waterbird abundance and Julian date (F_1, 789_ = 5.98, p = 0.015). This slowed, and became negative with time, as the quadric term of date (F_1, 788_ = 10.34, p = 0.0014) indicated a parabola of waterbird abundance within the migration season. There was also a significant interaction between year and the quadric term of date (F_1,768_ = 17.96, p < 0.001), showing that the parabola shape within the migration season changed across years. The average waterbird abundance (F_18, 790_ = 30.81, p < 0.001), and the annual trend (F_18, 769_ = 3.59, p < 0.001) differed between survey sites (Fig.S8a). The goodness of fit of the GLM was 0.511.

**Figure 5.**
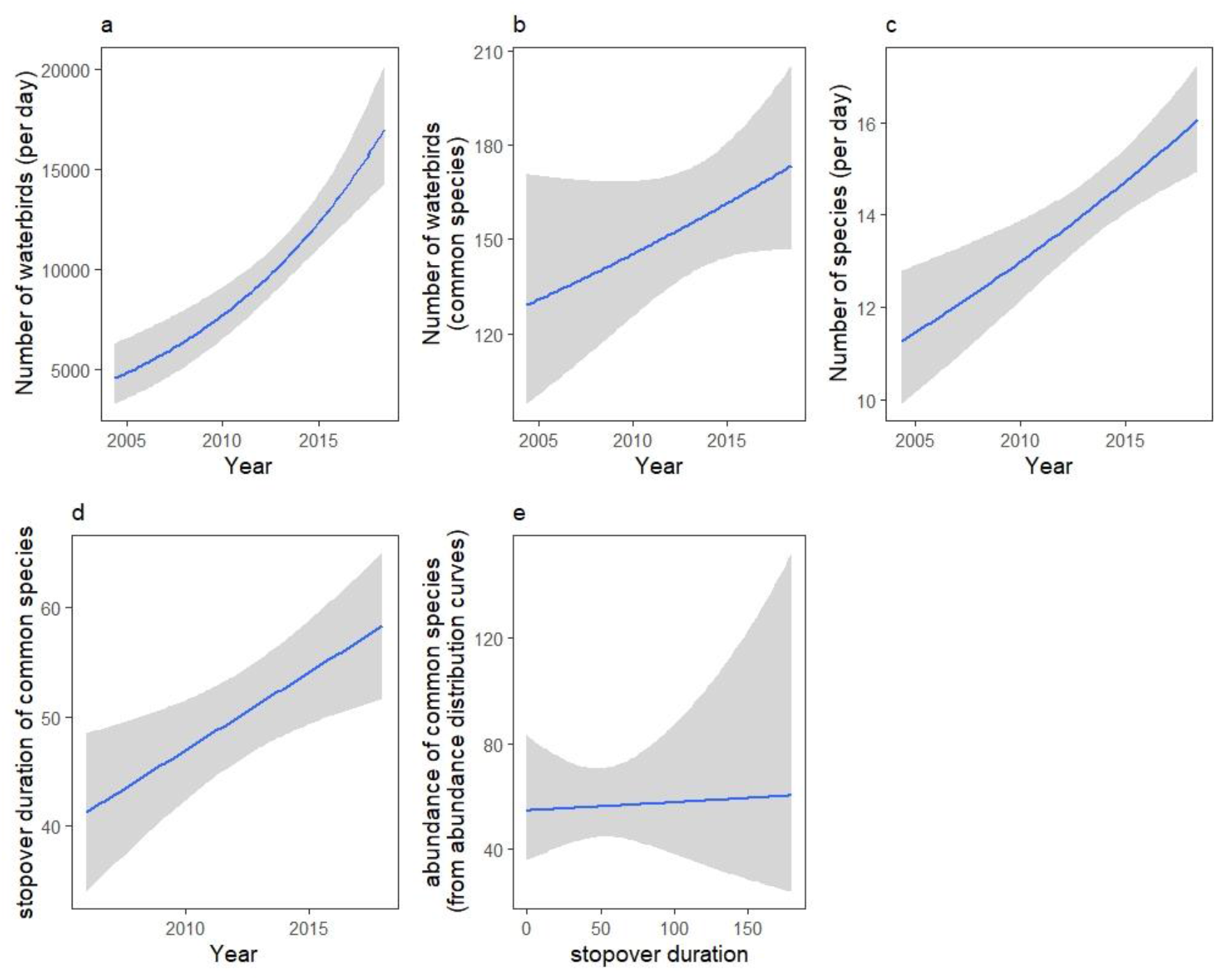
Trends of waterbirds abundance, richness and stopover duration from the census data. a) Overall waterbirds abundance as a function of year; b) the abundance of the most common species as a function of year; c) waterbirds richness as a function of year; d) stopover duration of common species as a function of year; e) The correlation between stopover duration and the abundance in common species

### Empirical data: the abundance of the most common species

In the analysis of the most common species, the abundance of common species increased at a rate of 35.13% per year (F_1, 18418_ = 6.92, p = 0.009) (Fig.5b), after correcting for effects from observers effort (F_1, 18417_ = 47.63, p < 0.001) and the interaction between year and effort (F_1,18372_=6.94, p=0.008), survey sites (F_18,18399_ = 95.28, p< 0.001) and the interaction between year and survey sites (F_18,18354_ =5.96, p< 0.001) (Fig. S8b), species (F_24,18373_ = 80.14, p < 0.001) and the interaction between year and species (F_24,18328_ = 6.96, p < 0.001). There was a quadratic association between Julian date and the number of birds (linear term, F_1, 18398_ = 25.77, p < 0.001; quadratic term F_1, 18397_ = 85.23, p < 0.001), but the interaction between year and Julian date was not significant (linear term, F_1, 18352_ = 0.52, p = 0.470; quadratic term F_1, 18353_ = 2.57, p = 0.109). The goodness of fit of the GLM was 0.443.

### Empirical data: waterbird richness

Waterbird richness significantly increased with time (F_1, 805_ = 28.20, p < 0.001) having corrected for effort (Fig.5c), with a rate of 7.93% per year. Effort was not statistically significant on richness (F_1, 804_ = 0.26, p = 0.61), although a weak interaction between year and effort was observed (F_1, 783_ = 4.69, p = 0.031). The relationship between richness and Julian date within a year was positive, with an initial rate of increase of 3.76% per day (F_1, 785_ = 34.50, p < 0.001). This rate slowed as Julian date increased, as the relationship between richness and the quadric term of Julian date was negative (F_1, 784_ = 63.55, p < 0.001), indicating an arched parabola in each migration season. The waterbird richness (F_18, 786_ = 29.16, p < 0.001) and yearly trend (F_18, 765_ = 4.69, p < 0.001) differed among survey sites (Fig S8c). The goodness of fit was 0.487.

### Empirical data: stopover duration

Analysis of the stopover duration of common species revealed a significant increasing trend at the rate of 1.39 days per year (F_1,245_ = 14.58, p < 0.001) (Fig.5d). The stopover duration (F_24,221_ = 7.20, p < 0.001) and its temporal trends (F_24,197_ = 1.90, p = 0.009) were differed among species (Fig S8d). The adjust R-square of the linear model was 0.428.

In the analysis of the correlation between bird abundance and the stopover duration, abundance was positively related with stopover duration (F_1,220_ = 18.36, p < 0.001) (Fig.5e), and also positively related with year (F_1,245_ = 56.41, p<0.001). The abundance (F_24,221_ = 35.93, p < 0.001) was differed across species. The relation between year and abundance (F_24,196_ = 3.28, p < 0.001), and the relation between stopover duration and abundance (F_24, 172_ = 2.24, p = 0.002) was differed across species. The interaction between year and stopover duration was not significant (F_1,171_= 0.70, p = 0.404). The goodness of fit of the GLM was 0.857.

## Discussion

By building a full-annual-cycle IBM of a stylized migratory population, we identify the critical role of stopover stage of northward migration in the influence of migration strategies and population dynamics of migratory birds across the whole annual cycle, and also identify the key processes linking individual migration strategy and population dynamics (Fig 6). Our empirical data provides evidence to support the mechanisms we showed from our theoretical model. Specifically, our results are consistent with the loss of staging habitat generating plasticity in migration strategy via increased intraspecific competition during migration stopovers. As the size of the staging area is reduced, individuals need to remain in the staging area for longer to obtain sufficient food to continue on their way due to an increase in the density of competitors. Our model shows that the consequence of this is individuals depart later, and often in poor condition, and fewer individuals make it to the breeding area. However, those that do make it fare well. The dynamics at the staging area can consequently have knock on effects on populations in the overwintering and breeding areas that impact the population dynamics across the annual cycle, by altering the component of the life history where population dynamics are regulated.

**Figure 6.**
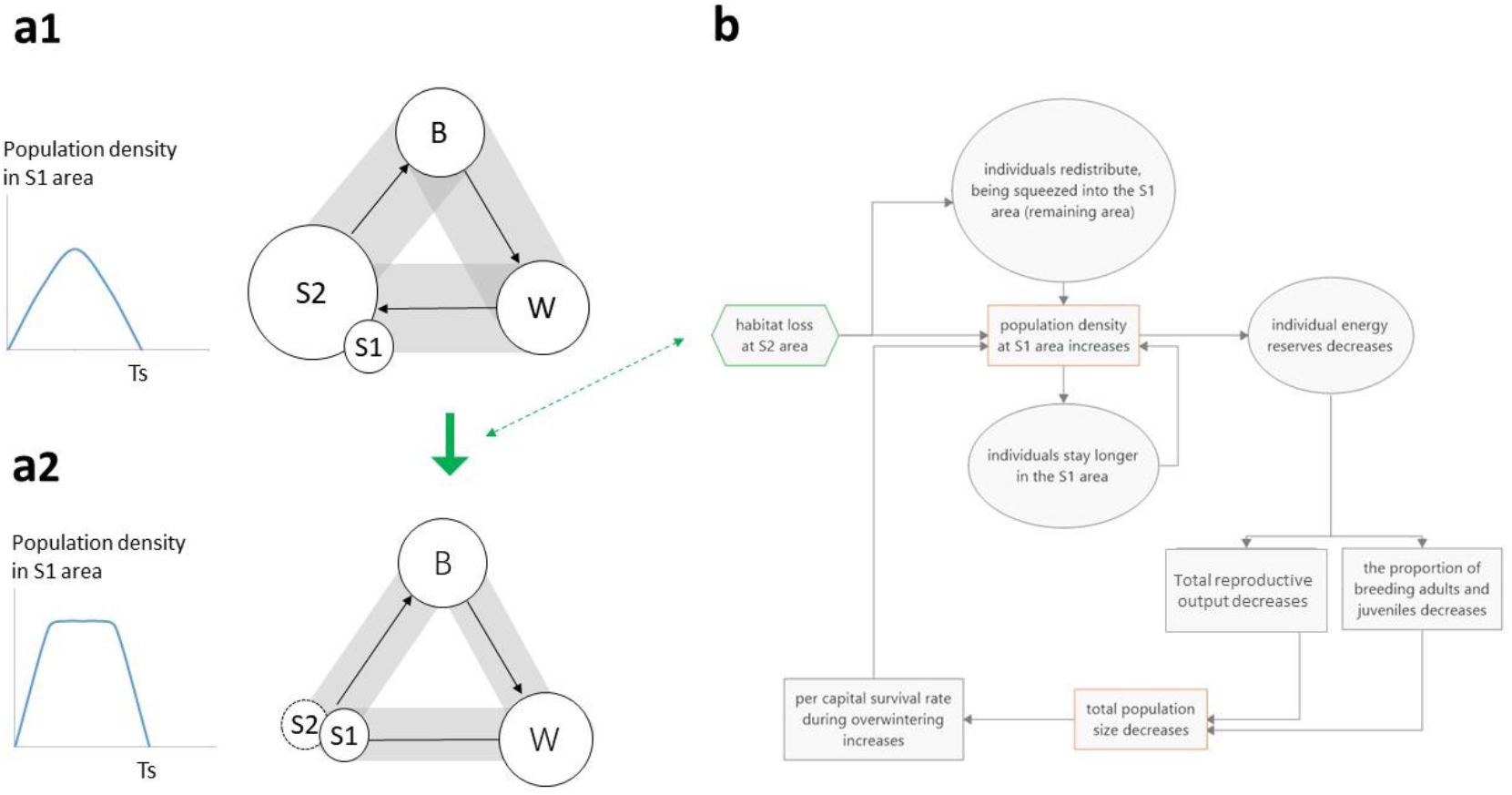
The processes linking individual migration strategy and population dynamics in our study. a) The changes in carrying capacity and population density as a function of the size of the staging habitat. S1 and S2 represents the two areas of the staging habitat, B represents the breeding habitat, and W represents the wintering habitat. The circle sizes represent the carrying capacity of each habitat, with the gray rectangles representing maximum population size across the annual cycle. In (a1) the size of the wintering and breeding habitats determine carrying capacity, while in (a2) the size of the staging habitat determines it. The line charts shows how population density in the staging areas changes during the period of migration stopover for scenarios (a1) and (a2) respectively. b) The feedback process between individual stopover duration and population dynamics.

Habitat loss in the staging habitat reduces the carrying capacity of the flyway, leading to population declines along it. As a consequence, the part of the annual cycle that determines the carrying capacity has been switched from the breeding habitat and wintering habitat to the staging habitat, as the spatial extent of staging habitat decreases. The staging habitat in the course of northward migration becomes the stage where the strongest density-dependence operates, and the population becomes regulated by the carrying capacity of staging habitat (Fig 6a). In contrast, competition on the breeding or wintering habitat is reduced, with processes operating in these areas no longer playing a major role in regulating the population dynamics. The contributions of different life history stages to the population dynamics can consequently vary with the spatial extent of the staging area. Such changes have the potential to alter selection on traits associated with competition for resources, and the entire life history.

Both in our simulations and empirical analysis, population density changes spatio-temporally following habitat loss in the staging habitat, with a positive correlation between bird numbers and stopover duration (Fig. 1b, Fig. S9 in Appendix S1). This contrasts the “buffer effect” hypothesis, which proposes that population density changes only in space (Brown 1969, Sutherland 1996a, Gill et al. 2001). Our results show that in addition to birds becoming more concentrated in the remaining area, a decrease in the extent of the staging area can also result in an increase in time spent there, leading to intensified competition during staging period. Therefore, during the time-limited northward migration, the reduced staging area not only leads to a higher population density in the staging habitat as individual birds stay longer, but also that the high population density is maintained for a longer period of time during this part of the life cycle (Fig.6a). And this can alter the strength of population regulation in either the wintering or breeding areas (Fig. 6b). How do we reach this conclusions?

In our models, individuals adjust their migration strategy to prolong the stopover duration when food becomes scarce. As a consequence, individuals often arrive at the breeding ground late and with low energy reserves, or they fail to reach the breeding grounds. This results in fewer adults arriving in the breeding ground, fewer adults reproducing, and a decline in total reproductive output. However, those individuals that do arrive experience less competition, and have higher departure energy reserves at the breeding habitat, breeding adults have higher per capita reproductive rate. All individuals have higher survival rate during southward migration and northward migration. Previous studies have reported evidence that support parts of the processes we describe here including: longer stopover duration is related to habitat loss, scarce food or high density of competitors (Moore and Yong 1991, Kelly et al. 2002, Conklin et al. 2021), decreased refuelling rate causes poor departure body mass in the staging area (Baker et al. 2004), limited refuelling time reducing survival rates in the northward migration (Rakhimberdiev et al. 2018), and reductions in total population-level reproductive output (Newton 2006, Desprez et al. 2018).

All of this is due to increased competition in the staging habitat, which reduces per capita food availability, and generates behavioural plasticity in migratory strategy. As individual behaviour changes, so to do the population dynamics. The altered population dynamics, in turn, affect individual behaviour (Miner et al. 2005). This process continues until an equilibrium is reached, when the behaviour settles down to equilibrium, as do the population dynamics (Fig. 6b). The connection between individual phenotypic plasticity and population dynamics is the result of feedback process across the annual cycle that can result in eco-evolutionary feedbacks are argued by Coulson (2020).

If the part of the annual cycle that determines carrying capacity is not switched to staging habitat, patterns and processes of individual strategies and population dynamics can be different (Fig. 4). The staging habitat was no longer the stage regulating population dynamics, the population density at the staging habitat no longer affected the trade-off between time and energy budgets in the course of stopover period, then influencing the life history process. The strongest density-dependent effects at the breeding habitat or the wintering habitat can cause a decrease in reproduction rate or survival rate at either stage, leading to a lower population density at the staging habitat, and a shortened stopover duration. Evidence has been reported by (Holmes et al. 1996, Rockwell et al. 2017) who observed reduced survival rates when the wintering habitat is limiting, and (Marra and Holmes 2001, Rodenhouse et al. 2003, Tomotani et al. 2018) who reported poor physical condition of juveniles and decreased reproduction when the breeding habitat is limiting. In addition, even though different breeding strategies may influence individuals’ energy budgets across the life cycle stages, our simulation results from model 2 and model 5 showed similar patterns and processes, indicating both capital breeders and income breeders would increase their stopover duration when facing habitat loss at staging habitats. Therefore, whether the changes in the extent of the staging habitat alters the part of the life cycle where carrying capacity is lowest will determine the direction of change in life-history processes. Our proposed process can only occur when the staging habitat become the stage that determine the carrying capacity of the whole annual cycle.

Our empirical and theoretical results align well, and are consistent with, patterns reported in the existing literature on the EAAF migration flyway, which are decreasing trends of total population size along the flyway, increasing trends of population density (Wilson et al. 2011, Yang et al. 2011, Clemens et al. 2016, Piersma et al. 2016, Studds et al. 2017) and increasing stopover duration at the remaining staging area(Conklin et al. 2021). Our study reveals the underlying mechanisms behind a seemingly positive phenomena observed in migratory birds, while also highlighting the crucial role of one life history stage to population dynamics across the whole annual cycle.

We call for further investigation on the connection between individual timing, energy reserves and demography across the whole annual cycle. Migratory birds need to be studied in depth at their breeding, wintering and staging areas to allow us to fully understand their dynamics. Focusing on a single area in detail, with limited data elsewhere, makes dynamical inference challenging, and may also lead researchers to reach erroneous conclusions. In addition, migratory species play a key role for a number of other ecological processes by transporting energy and nutrients between spatially distinct areas, or via their impacts on the ecological networks they form with other species in geographically separated areas (Bauer and Hoye 2014), understanding their dynamics is also important to further extend our knowledge on their roles in ecosystem stability. Although logistically difficult, and costly, our work suggests that studies of migratory species across the entire annual cycle are necessary if we are going to understand the dynamics of species that exhibit one of the most remarkable behaviours in the animal kingdom.

## Acknowledgements

We thank Hongyan Yang, Pinjia Que, Yajing Chang, Bingrun Zhu, Jingsheng Ma, Jianmin Wang, Paul Holt, Adrian Boyle and the Global Flyway Network team, and many volunteers for help with the fieldwork. This work was supported by the grants from the Natural Science Foundation of China (31830089 and 31572288 to ZZ, 31801985 to WL), the Paulson Institute grant and the SEE Foundation grant to ZZ, the China Scholarship Council grant to JL, the Fundamental Research Funds for the Central Universities to WL. CJH has been funded by GFN with grants secured by Theunis Piersma via BirdLife Netherlands (2007-2012), WWF Netherlands (2010-2014, 2016, 2018), Spinoza Premium of Netherlands Organisation Prize for Scientific Research to Theunis Piersma (2014-2017) and MAVA (foundation pour la nature) in 2018.

## Conflict of interest

None of the authors have a conflict of interest.

## Authors’ contributions

JL, TC and ZZ conceived the ideas and designed methodology. JL, WL, XM, and CJH collected the data. JL built the model, analysed the data and led the writing of the manuscript with significant contributions from TC. All authors contributed critically to the drafts and gave final approval for publication.

## Data availability statement

Code and data will be archived if the paper is accepted for publication.

## Notes

### Competing Interest Statement

The authors have declared no competing interest.

### Summary of Updates

New analyses have been added in both the modelling part and the empirical part

